# Prenatal exposure to particulate matter (PM_2.5_) from biomass fuel and low birth weight in a Sri Lankan birth cohort

**DOI:** 10.1101/461632

**Authors:** Meghan Tipre, Rajitha Wickremesinghe, Sumal Nandasena, Anuradhini Kasturiratne, Rodney Larson, Sreelatha Meleth, Udaya Wimalasiri, Claudiu Lungu, Tamika Smith, Nalini Sathiakumar

## Abstract

**Background:** Prenatal exposure to particulate matter of size 2.5 μm (PM_2.5_) emissions from biomass fuel may be associated with low birth weight (LBW) (<2.5 kg). We present results examining the association between PM_2.5_ and LBW from a birth cohort of 545 pregnant women followed from first trimester until delivery, in Sri Lanka.

**Methods:** Exposure to household air pollution (HAP) from biomass smoke was assessed using a detailed questionnaire; two-hour measurements of kitchen PM_2.5_ were collected in a subset of households (n, 303, 56%). Data from questionnaire and measured PM_2.5_ were used to estimate two-hour kitchen PM_2.5_ concentrations in unmeasured households. Fetal growth including weight and gestational age were assessed at birth. Linear and logistic regression were used to evaluate the association between HAP and birth weight.

**Results:** About 78% of the households used wood (n, 423). In linear regression models, we found an inverse association between a 10-unit increase in PM_2.5_ and birth weight (β±SE, −0.03±0.02; p, 0.02) adjusted for covariates. Similarly, households primarily using wood (>50%) were negatively associated with birth weight as compared to LPG users (β, −0.13±0.06; p, 0.03). In logistic regression models, a 10-unit increase in PM_2.5_ was associated with increased odds for LBW (OR, 1.26; 95%CI, 1.02-1.55; p, 0.04). The odds for LBW were highest among >50% wood users (OR, 2.82; 95%CI, 1.18-6.73; p, 0.01), compared to those using >50% LPG with wood (OR, 1.44; 95%CI, 0.57-3.63) and 100% LPG users (referent). The association between HAP exposure and birth weight/LBW were consistent among term-births (n,486).

**Discussion:** The finding of a significant association between prenatal PM_2.5_ exposure and LBW in a low-middle income country (LMIC) setting where competing risk factors are minimal fills a gap in the body of evidence linking HAP to LBW and underscore need to prevent and reduce of HAP exposure in LMICs.

## INTRODUCTION AND AIMS

Household air pollution (HAP) from combustion of solid fuels such as wood, dung and coal, used for cooking and heating, is ranked 10^th^ as a contributor to the global burden of disease (1,2) and the third leading risk factor for global mortality and disability-adjusted life years (DALYs)(3). Smoke emissions from solid fuel combustion in inefficient cookstoves and inadequate ventilation release relatively high concentrations of PM_2.5_, carbon monoxide (CO) and several other organic and inorganic compounds (4). Long-term exposure to fine particulate matter (PM_2.5_) of size 2.5 μm and smaller contributes to 4.2 million deaths and a loss of 103.1 million DALYs; majority of this burden is borne by low middle income countries (LMICs) (5)(6). More than three billion people accounting for 43% of the world’s population use solid fuels (coal and biomass) for cooking and heating; about 80% of these people live in LMICs (7,8). In most LMICs, women of child-bearing age typically spend several hours a day in cooking chores. Consequently, they are exposed to high levels of HAP for long periods and are at high risk for adverse health effects from HAP. In Sri Lanka, a LMIC in South Asia with a population of 20 million, about 74% of the population uses some form of solid fuel (unprocessed wood) for cooking.(8,9) Recent reports from the Sri Lankan National Demographic Surveys indicate that the prevalence of solid fuel use remains mostly unchanged with only slight decrease from 78% to 74% in the last 15 years. (10–12)

Studies on exposure to PM_2.5_ from either ambient or indoor sources have found positive associations with adverse pregnancy outcomes including stillbirths, low birth weight (LBW), small for gestational age (SGA), preterm births and birth defects (13–15). Results from a meta-analysis of ambient air pollution and birth weight found that birth weight in grams (g) was negatively associated with 10 μg/m^3^ increase in ambient PM_10_ and PM_2.5_ exposure during the prenatal period, adjusted for maternal smoking (16). Studies that have evaluated the role of HAP from solid fuel combustion and LBW indicate a stronger association, than ambient air exposure studies (17–20). A meta-analysis found that exposure to HAP from solid fuels based mostly on self-reports, resulted in an 86.4 g reduction in birth weight, and a 35% and 29% increased risk of LBW and stillbirth, respectively (21). Several of the pollutants found in ambient and indoor air share similar toxic properties as those found in environmental tobacco smoke (ETS) (22). Evidence from ETS and animal studies indicate that these air pollutants are absorbed by maternal blood, and then may cross the placental barrier and cause adverse effects on the fetus (23,24).

Previous studies of HAP and adverse birth outcomes including birth weight and LBW have several limitations. Most studies are either cross-sectional or case-control. Many of these studies are also subject to exposure misclassification. The labor-intensive and high costs associated with conducting air quality monitoring in all households, limit several HAP studies to qualitative exposure assessment resulting in exposure misclassification, especially in households with stacking of fuel (25). Stacking of fuel is where the household supplements the main fuel with secondary fuel for cooking certain dishes such as rice or beans or for special occasion such as festivals or holidays.

Most studies to date have been conducted in LMICs including India, Guatemala, Pakistan, and Ghana, countries where other competing risk factors for adverse birth outcomes such as malnutrition, poverty, infections and poor access to health care, are highly prevalent (21,26). Further, in-home deliveries and limited access to prenatal care in rural regions may often result in gaps in information related to fetal growth and pregnancy-related complications making it difficult to control for residual confounding factors to estimate the true effect of HAP on fetal health. Only a few were birth cohort or experimental in design but except for a study conducted by Balakrishna et. al., (2018), most had smaller sample size (27).

To address some of these challenges, we designed a prospective birth-cohort study in Sri Lanka. The goal of the study was to evaluate the association between prenatal exposure to HAP from biomass smoke emissions and adverse birth outcomes. Our choice of the study setting is unique and offered several advantages. Sri Lanka has a literacy rate of 92% and maternal-child health indicators comparable to most high-income countries (28); thus, confounding/competing risk factors are minimized. Next, Sri Lanka’s primary care infrastructure includes routine follow-up of almost all (>95%) pregnant women and their children until five years of age facilitating assembling a cohort of women early in their pregnancy, following them through childbirth, and to subsequently follow their children until five years of age.

## MATERIALS AND METHODS

### Study setting

Sri Lanka is divided into approximately 334 Medical Officers of Health (MoH) units areas under the management of a Ministry of Health; each MoH serves a population of 60,000 to 100,000 (29–31). The MoH area is further divided into several Public Health Inspector (PHI) areas and, within each PHI areas are Public Health Midwife (PHM) areas, the smallest administrative unit of the government health system in Sri Lanka. Typically, a PHI area serves a population of 10,000-15,000 which is subdivided into 3-4 PHM areas. The PHM is the female grass roots level family health worker primarily responsible for maternal-child health services and serving a population of 3000-5000 people (31,32).

The PHM registers newly pregnant women, typically before 10 weeks of gestation. Thereafter, women are followed at the field clinic and in their homes every four weeks until the 28th week of gestation, every two weeks from the 28th to the 36th week of gestation and weekly thereafter until delivery. The PHM is also responsible for mothers’ immunizations and health education. Almost 99% of births in Sri Lanka occur in hospitals. After delivery, the PHM visits the mother at home (a minimum of three visits within the first 10 days) to follow-up and identify early post-partum complications of the mother, if any, and newborn problems including feeding complications.

### Study area and recruitment

The study was conducted in the Ragama, Medical Officer of Health (MOH) which comprises of 16 PHM areas. Two research assistants (RAs) worked with the PHMs to identify eligible study participants. A total of 721 pregnant women (*n* = 721) were screened between July 2011 and June 2012 from the Ragama MOH area. Eligibility criteria included pregnant women in their first trimester; between 18 and 40 years of age; who planned to continue prenatal care and deliver at the Colombo North Teaching Hospital, Ragama; and permanently resided in the Ragama MoH area. Written informed consent in Sinhala was obtained from all women and the study procedures were explained prior to enrollment. Of the 721, 545 women were followed until delivery. Loss of participants were due to miscarriages (n, 60); twin births (n, 5); stillbirth (n, 9); delivered in different hospitals (n,13); lost to contact due to changed residence (n, 15) or refused to participate (n, 74).

### Data collection

Women were interviewed at enrollment (mean, 11±2.5 weeks of gestation) to collect baseline information on demographic factors, residential history, medical history, maternal and paternal occupational history, history of active and passive tobacco smoking; and household characteristics. A section of the questionnaire collected detailed exposure information on factors related to HAP including fuel use, kitchen characteristics and cooking practices. Information was also collected on other sources of indoor and outdoor pollution contributing to HAP. Maternal pregnancy history and infant anthropometric measurement at birth were obtained from hospital records. All interviews were conducted by trained bilingual study interviewers.

### Prenatal exposure assessment

Qualitative information on exposure to HAP was obtained from the baseline questionnaire administered to all study participants. The baseline questionnaire elicited detailed information on cooking practices such as types of fuels used for cooking including primary and secondary cooking fuels; percent use of each fuel during cooking; daily average hours spent cooking; type of stove; presence of a chimney and its condition; ventilation in the kitchen during cooking including number and size of open doors and windows; and the type of kitchen, its dimensions and relative position with other rooms. Information was also collected on other sources of HAP such as tobacco smoking habits of household members, burning of incense and mosquito coils, and proximity of sources of ambient air pollution such as vehicular traffic, industry or garbage disposal sites.

Quantitative exposure to HAP was ascertained from two-hour PM_2.5_ measurements in the kitchen during the cooking period in a subset of households (n, 304 out of 545). Based on baseline data, households were categorized into ‘high’ and ‘low’ exposure categories if the women used biomass fuel or clean fuel as primary cooking fuel, respectively. Using a stratified random sampling method, we selected 304 households (high exposure group, n, 158; low exposure group, n, 145) for conducting two-hour air quality measurements in the kitchen. At least one measurement per household was scheduled during the pregnancy period; in 106 households (36%), two sets of measurements were conducted in two of the three trimesters and in 26 households, three measurements were conducted, once every trimester; all others had one measurement (n,197, 65%). All measurements were conducted during a two-hour cooking session while preparing a typical lunch. Using this data, PM_2.5_ concentrations were estimated for unmeasured household (described below). PM_2.5_ concentrations were measured using a TSI Dusttrak II Aerosol Monitor 8530. Inlets of monitors were fixed at 145 cm above the floor, 100 cm from the cook stove and at least 150 cm away from an open window or door.

### Birth outcome assessment

All births occurred in the North Colombo Teaching Hospital. RAs collected data on birth outcomes from hospital records. Information abstracted included: birth weight, length and head circumference measured within 24 hours of birth, prenatal and delivery medical history. Gestational age was estimated based on the date of the last menstrual period and substantiated with ultrasound in the first trimester to classify infants as preterm vs. term births; pre-term births were defined as <37 weeks of gestation. LBW was defined as birth weight <2500 grams at birth. Term LBW were defined as full-term babies weighing <2500 grams at birth, as per standard definitions by the World Health Organization (33,34).

### Covariates and confounding variables

Data on covariates obtained from baseline questionnaire included: (i) demographic and SES factors such as maternal age, education, occupation, household income, type of housing, toilets, drinking water supply, and ownership of house; (ii) maternal lifestyle factors such as use of alcohol and tobacco; (iii) maternal exposure to toxic chemicals such as ETS, pesticides and lead; (iv) parity; (v) past and present obstetric history of pregnancy induced hypertension, gestational diabetes, other pregnancy-related complications, miscarriages and still births, iron deficiency, and calcium intake; (vi) and child’s gender.

### Ethical review

All study protocols were approved by the Ethics Review Committee of the Faculty of Medicine, University of Kelaniya and the Institutional Review Board of the University of Alabama at Birmingham (UAB).

### Statistical analysis

#### (i) Exposure estimation

##### Categorical HAP exposure

In the baseline questionnaire, information was collected on the primary fuel and supplementary fuels used in the households. A Likert scale was used to quantify the amount of fuels used typically during an average week. The scale ranged from 1 to 4, 1 being 0 to 25% of fuel use and 4 being 75% to 100% of fuel used. Most of the households (n, 423, 78%) in the study used a second fuel in varying quantities to supplement their primary fuel use. Combining the above information, we created two exposure variables. The first was a dichotomous variable classifying household into two categories: ‘only LPG’ and ‘any wood use’. The second variable classified the households into three categories with gradients of wood use: (i) those using >50% to 100% of wood; (ii) those using >50-100% of LPG or kerosene and supplementing it with <50% wood; and (iii) those using 100% LPG. Since, only three households used 100% kerosene supplemented occasionally by LPG, these households were included in the third fuel category of 100% LPG.

##### Continuous PM_2.5_

Next, using regression models, we examined the influence of several physical characteristics on concentration of PM_2.5_ levels in the kitchen fuel types (>50% wood, >50% LPG + <50% wood, 100% LPG), kitchen type (indoor, temporary hut or outdoor); cookstoves (traditional cookstove (TCS), improved cookstove (ICS), kerosene or LPG stove); presence or absence of functional chimney, number of doors and windows open at the time of cooking as a surrogate for kitchen ventilation, and average daily cooking time categorized into three groups (0 to 60 minutes; 60 to 90 minutes and >90 minutes).

###### Prediction model

Data from 275 out of 303 households with air quality measurements were used to develop the exposure model to estimate kitchen PM_2.5_ airborne concentrations. Twenty-six samples were excluded as the comparison between fuel use and PM_2.5_ values suggested potential misclassification of fuel use or were identified as possible outliers. For example, a household using >75% of LPG had measured value of PM_2.5_ as high as 1000μg/m^3^ significantly greater than the concentration that would be expected. The final regression model was then used to quantify kitchen for PM_2.5_ concentrations in kitchens without air quality measurements (n, 270; 50%); data on percentage of fuel use was missing for one subject and thus, had missing value for PM_2.5_.

Data (n, 275) was partitioned randomly into training (n, 192) and test (n, 83) datasets using a 7:3 ratio respectively. The training dataset was used to identify factors significantly associated with log-transformed PM_2.5_. Linear regression was used to model log-transformed PM_2.5_ as a function of significant risk factors. In univariate statistics, geometric means for PM_2.5_ were compared across physical variables. Factors significantly associated with log-transformed PM_2.5_ at α <0.05 were included in a multivariable linear regression model to derive the final model. Adjusted R^2^ was used to evaluate the model fit. Results identified three independent risk factors associated with log-transformed PM_2.5_ including multilevel fuel categories, kitchen type, and functional chimney at p <0.05 (Supplementary table 1-2). The model R^2^ was 0.57 (supplementary table 2). The regression model equation is described below.

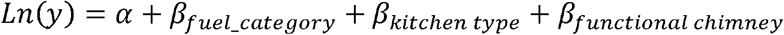

*where:* α, intercept =4.31; β_wood_ = 2.10, estimate for wood use >50%; β_LPG+Wood_ = 0.42, estimate for LPG use 50% to 100% supplemented by wood; β_LPG_ = 0, estimate for 100% LPG; β_K_ind_ = −0.18, estimate for indoor kitchen; β_K_temphut_ = 1.17, estimate for temporary outdoor hut kitchen; β_K_outdoor_ = 0, estimate for outdoor open kitchen; and β_chimney_ =0.06, estimate for functional chimney.

###### Validation of the model

In the test dataset, the beta (β) coefficients from training model were β used to predict PM_2.5_ values. The difference between the observed and the predicted PM_2.5_ values were calculated and residuals were plotted to evaluate the robustness of the predictive model (Supplementary table 2).

#### (ii) HAP exposure and outcomes

Initial analyses compared the mean distribution of birth weight and prevalence of LBW with exposure variables and covariates using t-test, ANOVA, and the chi-square test (supplementary table 1). We restricted the latter comparisons and further analyses to those potential risk factors, to which at least 10 subjects were exposed.

HAP exposure was analyzed as continuous (log-transformed PM_2.5_) and categorical variables (dichotomous and three-level fuel categories). We used linear regression to evaluate the association between HAP exposure and birth weight. Covariates significantly associated with exposure and/or birth weight in bivariate statistics at p<0.05 were included in the multivariable linear regression to compute adjusted beta-estimate and standard error, and adjusted means for birth weight.

Logistic regression was used to evaluate the association between HAP exposure and LBW. Covariates significantly associated with exposure and/or LBW in bivariate analysis at p<0.05 were included in the multivariable model. The odds ratio (OR) and 95% confidence intervals (CIs) were computed for LBW in relation to HAP exposure, modeled as continuous or categorical variable, adjusting for other potential risk factors.

We further stratified the analysis by term and preterm births The association between HAP exposure and birth weight were further examined stratified by term and preterm births. All analyses were conducted using SAS version 9.4 (SAS Institute NC).

## RESULTS

Of the total mothers (n, 545) followed-up from 1^st^ trimester until delivery, 423 (78%) used wood in varying quantities, while 122 (22%) women used only LPG for cooking. The average age of mothers enrolled was 29 years (SD, 4.9; median, 29); all women were married. Significant differences were noted between wood users and LPG users for maternal education, spouse’s education, main household water supply, ownership of the household, type of house (semi-permanent vs. permanent), income, kitchen type, functional chimney, and sources of HAP other than biomass fuel including ETS, burning of candles or incense (p<0.05). No remarkable differences were noted for mother’s age at the time of enrollment, past and current obstetric history, parity or child’s gender (Table 1).

**Table 1.**
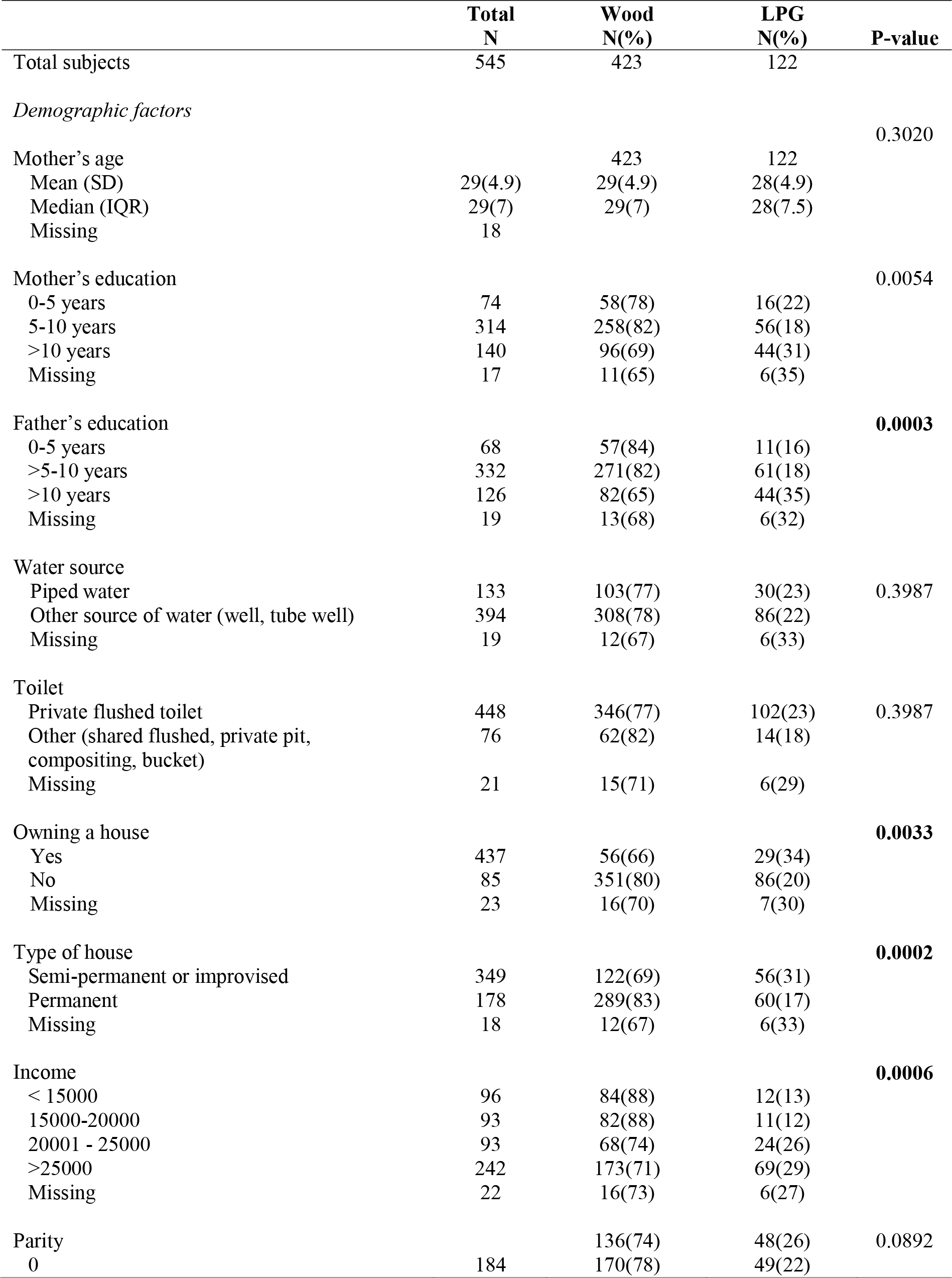
Descriptive statistics comparing variables between ever wood users and LPG users

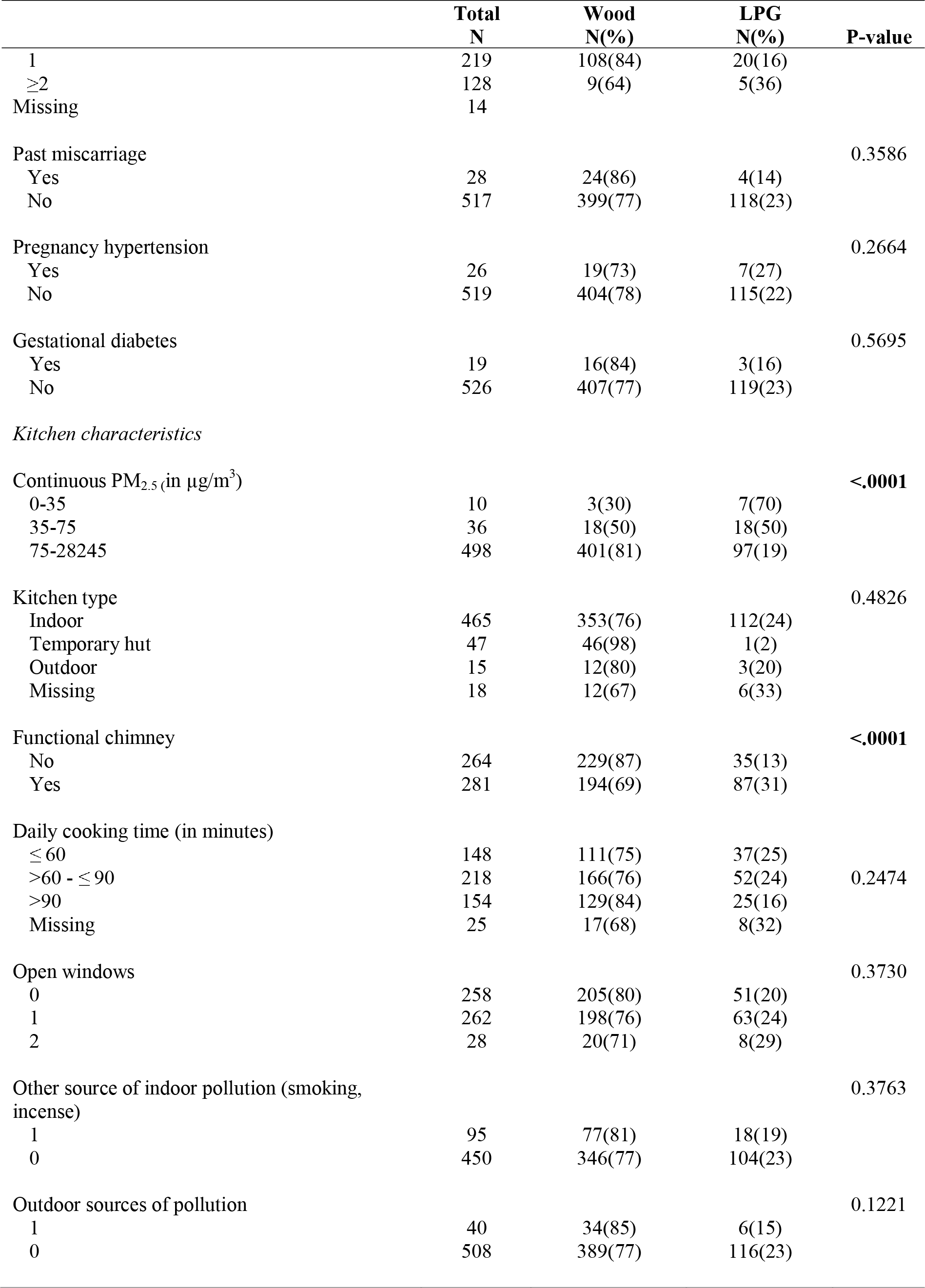

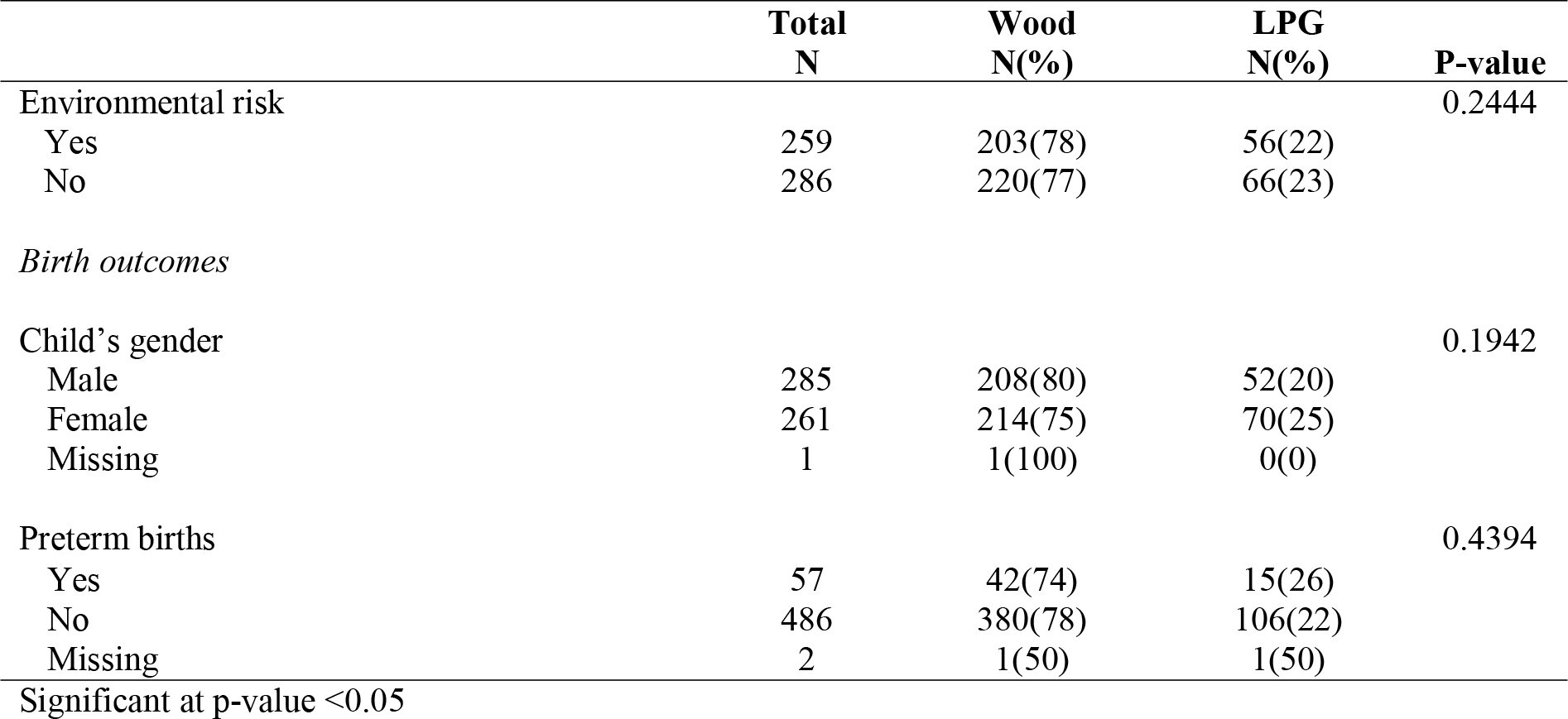

Among households with air quality measurements, the geometric mean ± standard error for PM_2.5_ among wood users (n, 157; 737± 80μg/m^3^) was about four times higher than households using >50% of LPG with wood (n, 78; 198±29 μg/m^3^) and almost 7 times higher than households using 100% LPG (n,68;100 ±10μg/m^3^).

### HAP exposure and birth weight

Of the total 545 newborns, 52% were boys. The mean birthweight of infants in the cohort was 2.95 (±0.45) kg. The proportion of LBW children in the cohort was 13% (n, 70 of 545) and the preterm births were 10% (n, 57). The total number of term LBW births were 46 (10%) out of 486 term births. About 65% of the mothers had vaginal delivery including forceps (n,6) and vacuum deliveries (n, 5).

Table 2 presents distribution of mean birth weight across all covariates. Birth weight was significantly associated with continuous PM_2.5_, three-level fuel categories, type of house, pregnancy induced hypertension and preterm births at p <0.05. In multivariable linear regression models (Table 3), continuous log-transformed PM_2.5_ was inversely associated with birth weight (β, −0.06; SE, 0.02; p, 0.03), adjusted for covariates. All multivariable models were adjusted for mother’s education, father’s education, income, permanent or semi-permanent house, ownership of the house, and pregnancy induced hypertension. In addition, models with categorical HAP exposure were also adjusted for type of kitchen and functional chimney.

**Table 2.** Distribution of mean birth weight and prevalence of LBW across demographic, maternal and household characteristics

|  | Total<br>N | Birth weight<br>Mean (SD) | LBW<br>N(%) |
| --- | --- | --- | --- |
| Total subjects | 545 |  | 70(100) |
| *Demographic factors* |  |  |  |
| Mother's age |  |  |  |
| Mean (SD) | 29(4.9) |  | 28(4.9) |
| Median (IQR) | 29(7) |  | 28(5.5) |
| Missing | 18 |  |  |
| Mother's education |  |  |  |
| 0-5 years | 74 | 2.9(0.5) | 9(12) |
| 5-10 years | 314 | 2.9(0.5) | 41(13) |
| >10 years | 140 | 3(0.5) | 18(13) |
| Missing | 17 |  | 2(13) |
| Father's education |  |  |  |
| 0-5 years | 68 | 2.9(0.5) | 12(18) |
| >5-10 years | 332 | 2.9(0.5) | 45(14) |
| >10 years | 126 | 3(0.5) | 11(8.7) |
| Missing | 19 |  | 2(11) |
| Water source |  |  |  |
| Piped water | 133 | 3(0.5) | 12(9.0) |
| Other source of water (well, tube well) | 394 | 2.9(0.5) | 56(14) |
| Missing | 19 |  | 2 (11) |
| Toilet |  |  |  |
| Private flushed toilet | 448 | 2.9(0.5) | 60(13) |
| Other (shared flushed, private pit, composting, bucket) | 76 | 3.0(0.5) | 8(11) |
| Missing | 21 |  | 2 (10) |
| Owning a house |  |  |  |
| Yes | 437 | 3.0(0.5) | 58(13) |
| No | 85 | 2.9(0.4) | 10(12) |
| Missing | 23 |  | 2 (8.7) |
| Type of house |  |  |  |
| Semi-permanent or improvised | 349 | **2.9(0.5)\*\*** | 51(15) |
| Permanent | 178 | **3(0.4)** | 17(10) |
| Missing | 18 |  | 2(11) |
| Income |  |  |  |
| < 15000 | 96 | 2.9(0.4) | 11(11) |
| 15000-20000 | 93 | 3(0.5) | 10(11) |
| 20001 - 25000 | 93 | 3(0.5) | 16(17) |
| >25000 | 242 | 2.9(0.5) | 30(12) |
| Missing | 22 |  | 3(14) |
| Parity |  |  |  |
| 0 | 184 | 2.9(0.4) | 23(13) |

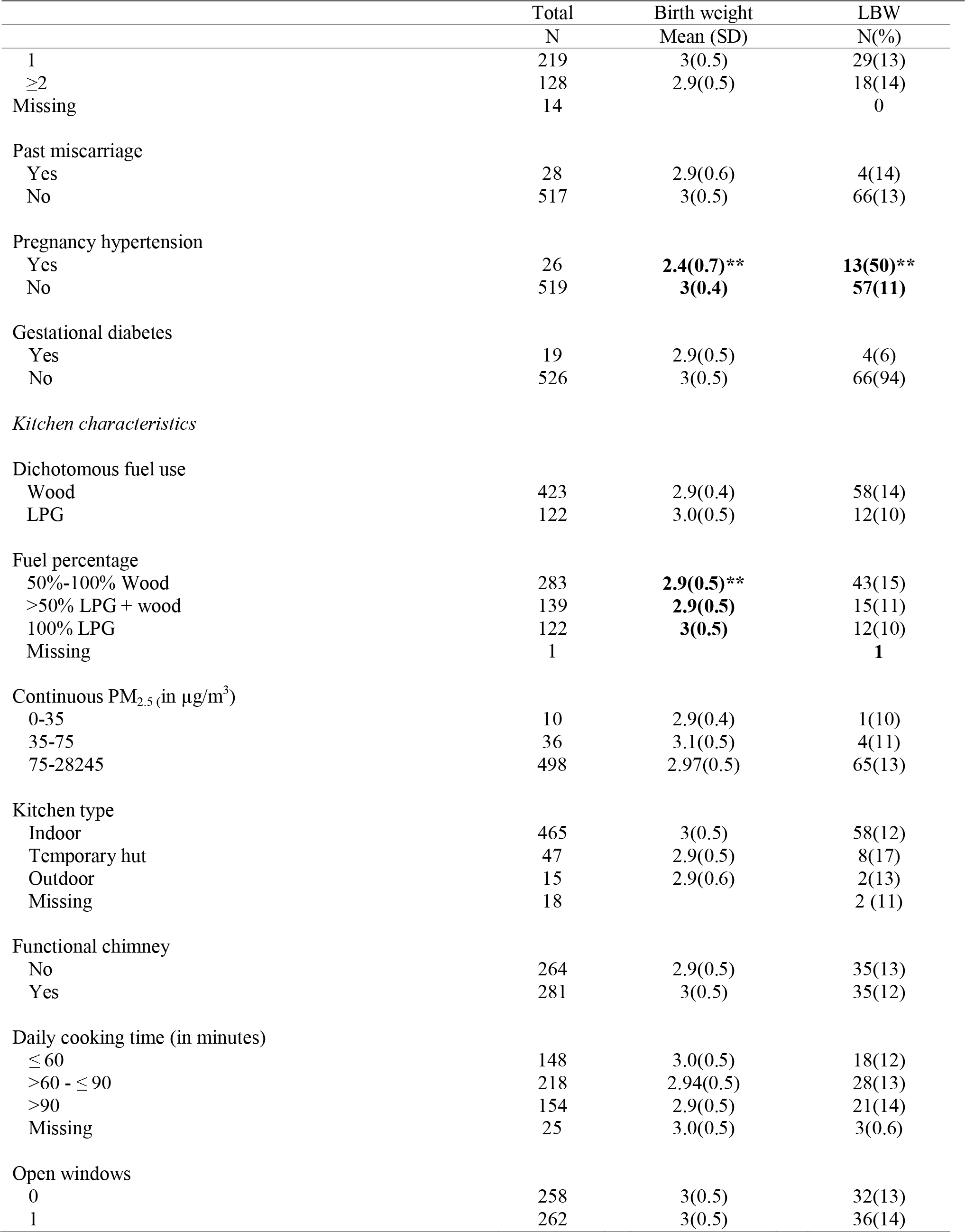

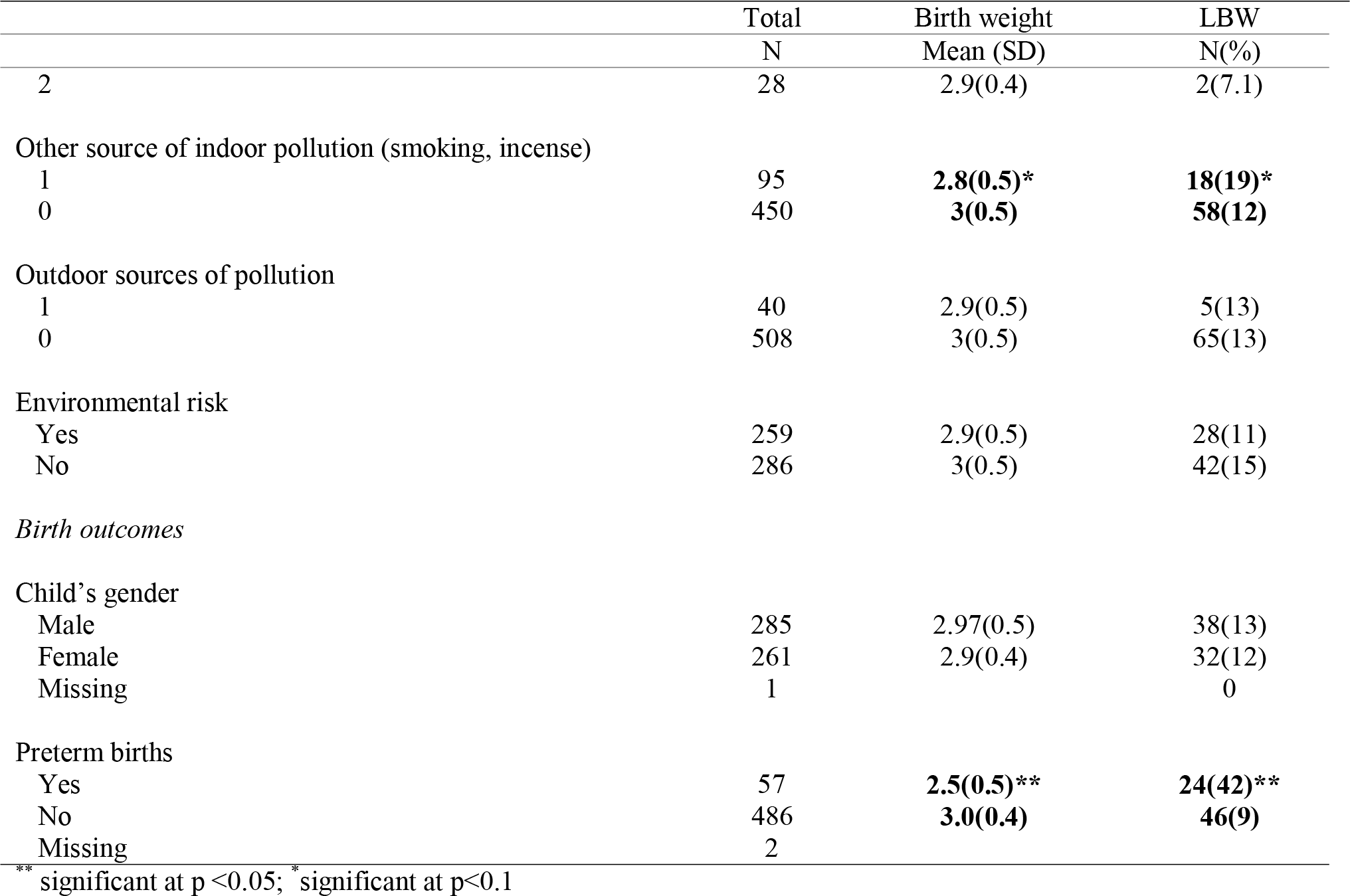

**Table 3.** Relationship between HAP exposure and birth weight (in kgs)

| | Exposure | Crude $\beta$<br>(Standard error<br>(SE)) | Adjusted $\beta$(SE) | P-value | Adjusted R<sup>2</sup>,<br>sample size |
| --- | --- | --- | --- | --- | --- |
| Model<sup>1</sup> | Continuous PM<sub>2.5</sub> | -0.06(-3.17) | **-0.03(0.02)** | **0.0339** | 0.10 |
| Model<sup>2</sup> | Wood | -0.07(0.05) | -0.07(0.05) | 0.2008 | 0.09 |
|  | LPG | 0 |  |  |  |
| Model<sup>3</sup> | >50% wood | -0.10 (0.05) | **-0.13(0.06)** | **0.0229** | 0.11 |
|  | >50% LPG + wood | 0.02 (0.06) | 0.002(0.06) | 0.9724 |  |
|  | 100% LPG | 0 |  |  |  |

<sup>1</sup>Adjusted for mother's education, father's education, type of house, ownership of the house, toilet, income, average daily cooking time, other sources of indoor pollution, pregnancy induced hypertension
<sup>2,3</sup>Adjusted for mother's education, father's education, type of house, ownership of the house, toilet, income, kitchen type, presence of functional chimney, average daily cooking time, other sources of indoor pollution, pregnancy induced hypertension

Use of wood fuel as compared to LPG use was associated with lower birth weight but the association was non-significant (β,−0.07; SE, 0.05; p, 0.20). In model with three-level HAP exposure, with >50% wood use (β, −0.13; SE, 0.06; p, 0.03) was significant and inversely associated with birth weight as compared to 100% LPG use (as referent). No association was observed between >50% LPG with wood use and birth weight (β, 0.002; SE, 0.06; p, 0.53).

### HAP exposure and LBW

The prevalence of LBW was significantly associated with continuous PM_2.5_; a 10 unit (in μg/m^3^) increase in PM_2.5_ was associated with 1.26 times increased odds of LBW (95% CI, 1.01-1.55; p, 0.04). No association was observed between dichotomous fuel use variable and LBW (OR, 1.78; 95% CI, 0.78-4.09; p, 0.17). For three-level HAP exposure variable, the odds of LBW were significantly higher for >50% wood users (OR, 2.27; 95% CI, 0.94-5.12; p, 0.08) compared to 100% LPG users; no difference was noted between >50% LPG with wood users and 100% LPG users (Table 4).

**Table 4.** Relationship between HAP exposure and LBW

|  | Exposure | Crude OR<br>(95% CI) | Adjusted OR (95%<br>CI) | P-value |
| --- | --- | --- | --- | --- |
| Model<sup>1</sup> | Continuous PM<sub>2.5</sub> | 1.14(0.94-1.38) | **1.24(1.01-1.55)** | **0.0428** |
| Model<sup>2</sup> | Wood | 1.46(0.76-2.81) | 1.82 (0.81-4.11) | 0.1500 |
|  | LPG | 1.00 | 1.00 |  |
| Model<sup>3</sup> | >50% wood | 1.64 (0.83-3.24) | **2.14(0.90-5.07)** | **0.0791** |
|  | >50% LPG + wood | 1.11 (0.50-2.47) | 1.44(0.57-3.63) | 0.6377 |
|  | 100% LPG | 1.00 | 1.00 |  |

<sup>1</sup>Adjusted for mother's education, father's education, type of house, ownership of the house, toilet, income, average daily cooking time, other sources of indoor pollution, pregnancy induced hypertension
<sup>2,3</sup>Adjusted for mother's education, father's education, type of house, ownership of the house, toilet, income, kitchen type, presence of functional chimney, average daily cooking time, other sources of indoor pollution, pregnancy induced hypertension

#### Term births

In stratified models by term birth, the relationship between exposure variables and birth weight and LBW were consistent as observed in the full models but effect estimates were lower. A moderately significant association was found between 10-unit increase in PM_2.5_ and decrements in birth weight (β, −0.04; SE, 0.02; p, 0.03) and LBW (OR, 1.27; 95% CI, 0.98-1.65, p, 0.07). Similarly, >50% wood use was associated with decrease in birth weight (β, −0.12; SE, 0.06; p, 0.05) and prevalence of LBW (OR, 1.76; 95% CI, 0.61-5.03; p, 0.08), but the associations were not significant. We did not run models on preterm birth due to small number of observations (n, 55).

## Discussion

In the present study, we investigated the relationship between prenatal exposure to HAP from biomass fuel use for cooking and birth weight, in a birth cohort study of 545 mother-child pairs in the Western Province of Sri Lanka. Our study modeled exposure to HAP as continuous and categorical variables. We found a significant association between two-hour kitchen concentrations of PM_2.5_ and birth weight and prevalence of LBW, after adjusting for significant confounding factors. Similarly, households using most wood were associated with decrease in birth weight and increased prevalence of LBW as compared to households using clean fuels. The results are consistent with previous air pollution studies including ambient and HAP and LBW (15,16,35). In our study, a combination of qualitative and quantitative data from household with measured PM_2.5_ were used to quantify kitchen concentrations of PM_2.5_ in unmeasured households. To our knowledge, our study is the first to use both measured and estimated PM_2.5_ measurements to evaluate the association between PM_2.5_ and birth weight. The exposure estimation method, effect estimates for birth weight and LBW in context of previous literature and the strengths and limitations of the study are discussed below.

### HAP, birth weight, LBW and previous studies

The results of our birth-cohort evaluating the relationship between HAP and LBW were consistent with the largest birth-cohort study to date as well as several meta-analyses published in the last decade. Balakrishnan et al. (2017) reported a 4 grams (95% CI, 1.08-6.76) decrease in birthweight and 2% increase in prevalence of low birthweight (OR, 1.02; 95% CI, 1.01-1.04) with a 10-μg/m^3^ increase in PM_2.5_. Results from other meta-analyses also found increased prevalence of LBW between 10% to 45%, associated with HAP exposure (17,19,21,36–40). Similarly, a recent meta-analysis conducted by Sun et al. (2016), found significant association between 10μg/m^3^ increment of ambient PM_2.5_ and increased odds of LBW (OR,1.09; 95% CI, 1.03-1.15) (35). The effect estimates in our study are closer to the higher end of central estimates reported by individual studies and meta-analyses. In our study, the average two-hour kitchen PM_2.5_ in kitchens using wood was almost 12 times higher than households using LPG (1649 vs. 143). Balakrishnan et al., (2018) reported 24-h average kitchen concentrations in households which were much lower than those reported in our study (wood use: 229±233 μg/m^3^; vs. 1649±166 μg/m^3^; LPG/electricity: 59±46 μg/m^3^ vs. 143±12 μg/m^3^), respectively. The higher concentration of kitchen PM_2.5_ found in our study may account for the higher effect estimate reported in our study as compared to the results reported by Balakrishnan et.al. (2016). However, without additional data on 24-hour average kitchen concentration from our study, no firm conclusions can be drawn.

The increased odds for term LBW (OR = 1.69; 95% CI = 1.03, 2.02) associated with 10-unit increase in PM_2.5_ were moderately higher than those reported by Tielsch et al., (2009) (OR = 1.21, 95% CI= 1.11, 1.31)(41) and Mavalankar et al., (1992) (OR=1.23; 95% CI : 1.01, 1.49) (42).

### Exposure estimation

In our study, information on stacking of fuel during an average week was used, to categorize HAP exposure into three categories. We developed a predicted model using information on fuel stacking, type of kitchen, and chimney to predict kitchen PM_2.5_ concentrations. This novel approach was used to evaluate the feasibility of using questionnaire responses to predict PM_2.5_ concentration in kitchen during cooking period, for households that did not have air pollution measurement data. Cross-validation approach evaluated the robustness of the model with a R^2^ of 57%. Personal monitoring of exposure to HAP by wearing portable devices for a period of 24 to 48 hours can substantially reduce exposure misclassification and improve the power of the study to detect relationships between exposure to HAP and adverse health outcomes (43). However, conducting area or personal air quality monitoring in all study subjects significantly increases study costs and respondent burden.

Previous HAP studies have mostly relied on questionnaire responses as surrogate for exposure. In more recent studies, questionnaire data is supplemented with area measurements for short time periods (snapshots) at multiple time points to generate time-integrated exposure metric to characterize chronic exposure (25). In these studies, households are categorized based on their primary fuel use as households using wood, kerosene or LPG and electricity. Few studies have considered multiple types of fuel use or the proportion of fuels used, resulting in uncertainty in nature and magnitude of exposure variability and misclassification of exposure in epidemiologic studies (25).

Balakrishnan et al.(2013), conducted one of the first studies that included 24-hour PM_2.5_ measurements in all study households (n, 474). They fitted a regression analysis to model 24-hour data to predict household PM_2.5_ concentrations (44). They identified fuel type, kitchen type, ventilation, geographical location and cooking duration, to be significant predictors of PM_2.5_ concentrations (R^2^= 0.33) with a fair degree of correlation (r=0.56) between modeled and measured values.(45) Another study conducted by Baxter et al. (2007), used questionnaire information combined with ambient air quality measurements to predict indoor concentrations of nitrogen dioxide, PM_2.5_, and elemental carbon in lower socio-economic status homes in urban Boston, Massachusetts (46). Their study found that outdoor concentrations, cooking time and occupant density were significant predictors of indoor PM_2.5_ concentration (adjusted R^2^ =0.36). The inclusion of open windows as an effect modifier for ambient air increased the R^2^ to 0.40. A recent study conducted by Zhou et al. (2018) in Shanghai, China, used outdoor air measurements and household characteristics to predict outdoor PM_2.5_ infiltration of indoor air (47). The models stratified by seasons and including information on air conditioning use, meteorological factors, floor of residence and building age, predicted 60.0%-68.4% of the variance in 7-day averages of indoor PM_2.5_ infiltration by outdoor air.

The R^2^ for the exposure model developed in our study (57%) was lower than the R^2^ (66%) reported by Zhou et al (47); and 1.6 times greater than the R^2^ (0.36) reported by Balakrishnan et al. (2013) (45). The correlation between estimated and measured PM_2.5_ (r, 0.74 vs. r, 0.56) was also higher in our study compared to that reported by Balakrishnan (2013) study.

Our results indicate moderate feasibility of using detailed questionnaire information to quantify kitchen PM_2.5_ concentrations. Several other factors can affect personal exposure including the high degree of variability from stacking of fuels, different stove types and kitchen designs, and time-activity pattern of the subjects i.e., the amount of time spent in the kitchen and inside the homes as compared to outside. Furthermore, changes in patterns of fuel use over the course of pregnancy could change personal exposure. Increased frequency, longer duration of air quality monitoring and time activity diary of time spent indoor could improve the accuracy of our predictive model.

Our study has several strengths. HAP exposure was analyzed in several different ways with consistent results. The retention rate in the study was almost 84% (545/656) after accounting for miscarriages, stillbirths and twin pregnancies. The study participants were identified through routine prenatal checkups at the antenatal clinics and were representative of the study population. All births occurred in hospitals and birth weight was recorded within 24-hours. Gestational age was determined by ultrasound examination in 1^st^ trimester if there was doubt about the last menstrual period, consequently, reducing information bias for LBW. In Sri Lanka, government-funded public health system provides prenatal and antenatal care to almost 99% of the pregnant mothers. Thus, self-reported maternal medical history was verified with maternal pregnancy records obtained from the antenatal clinics. Maternal lifestyle, such as smoking cigarettes, and use of alcohol, are known confounding factors for adverse birth outcomes. However, none of the mothers in our study smoked or consumed alcohol. Information on ETS was self-reported but no significant differences were noted for LBW and normal children. We were able to collect and adjust for known confounding factors associated with LBW including pregnancy induced hypertension, gestational diabetes, and other complications during pregnancy such as essential hypertension, high head, hyperemesis, hyperthyroidism, hypothyroidism, and threatened miscarriage.

The major limitation of our study was the lack of 24-hour air quality measurements for indoor and ambient air quality. Data on indoor air quality was available for 60% of households in the study and were used to quantify indoor air quality for remaining 40%, which could have resulted in some exposure misclassification. However, internal consistency in results suggest minimum bias.

## Conclusion

Our study evaluating HAP and birth outcomes in Sri Lanka adds to the growing evidence linking HAP to LBW and term LBW. Although, air quality measurements were conducted in a subset of households, both qualitative and quantitative data from measured households were used to quantify HAP exposure in households without measured data. Results of this study are important as almost 74% of the population in Sri Lanka continue to use wood for cooking. Since almost one-third of world’s population is exposed to HAP from solid fuel, even modest increase in the risk of adverse pregnancy outcomes has large-scale implications on the overall health of the population.

## Acknowledgements

The Prenatal Exposure to Biomass Smoke and Infant Neurodevelopment in Sri Lanka, study was supported by the National Institutes of Environmental Sciences (NIEHS: grant R21ES018730).

