## Supplemental table 1 and 2 for "Prenatal exposure to particulate matter (PM_2.5_) from biomass fuel and low birth weight in a Sri Lankan birth cohort"

| Supplementary table 1. Descriptive statistics of exposure-related variables by data source, distribution of PM_2.5_ across covariate categories | | | |
| --- | --- | --- | --- |
| Household kitchen characteristics | N (%) | Arithmetic (SE) | Geometric (SE)^†^ |
| Total N | 275^*^ |  |  |
| Fuel use |  |  |  |
| Any wood use^†^ | 209(76) | 1290 (133) | 475 (46) |
| Only clean fuel use | 66(24) | 116 (11) | 100 (11) |
| Fuel use by type and percentage |  |  |  |
| >50% wood | 141(51) | 1620 (167) ** | 740 (81) |
| >50-100% LPG and <50% wood | 78(25) | 785 (368) | 193 (29) |
| 100% LPG, Kerosene or Electricity | 66(24) | 179 (48) | 100 (11) |
| Missing |  |  |  |
| Kitchen |  |  |  |
| Indoor | 249 (91) | 870 (94) ** | 298(27) |
| Temporary hut | 15 (5.6) | 3290 (993) | 1843(527) |
| Outdoor | 10(3.7) | 1085 (301) | 638(238) |
| Missing | 1 |  |  |
| Functional chimney |  |  |  |
| Yes | 147(53) | 1114(135) | 256(33) |
| No | 128(47) | 886.7(164) | 433(51) |
| Number of open windows during cooking |  |  |  |
| Missing |  |  |  |
| 0 | 105(38) | 1254(207) | 415(62) |
| 1 | 153(56) | 838(108) | 306(35) |
| 2 | 17(6.1) | 1020(574) | 246(95) |
| Other indoor pollutant sources^‡^ |  |  |  |
| Yes | 47(17) | 1062(251) | 401(82) |
| No | 228(83) | 997(116) | 327(33) |
| Immediate outdoor pollution^¥^ sources |  |  |  |
| Yes | 18(10) | 1006(242.7) | 441.3(106.5) |
| No | 257(90) | 1008(113.9) | 330.4(31.5) |
| Cooking time (in minutes) |  |  |  |
| 0-60 | 74 | 844 (212) ** | 257 (44) |
| 61-90 | 118 | 928 (128) | 330 (45) |
| >90 | 80 | 1303 (239) | 463 (75) |
| * Excludes subjects with extreme values and/or inconsistencies between primary fuel type and primary stove type.  ^‡^Including indoor cigarette smoking, burning of incense, mosquito coil, and candles inside the homes  ^¥^Includes industry near the homes  ^g^Average daily cooking time across 2-3 meals per day  **p<0.05; the av | | | |

| Supplementary table 2. Regression model to predict log-transformed PM_2.5_ kitchen concentrations during cooking periods | | | | | | | | | | |
| --- | --- | --- | --- | --- | --- | --- | --- | --- | --- | --- |
| Parameter |  | Estimate | Standard  Error | t Value |  | Cross Validation Estimates | | | | |
|  |  |  |  |  | P-value | 1 | 2 | 3 | 4 | 5 |
| Intercept |  | 4.72 | 0.44 | 9.13 | <.0001 | 4.48 | 4.54 | 4.80 | 4.78 | 4.86 |
| Fuel categories |  |  |  |  |  |  |  |  |  |  |
| 50%-100% Wood |  | 2.14 | 0.19 | 2.12 | <.0001 | 2.15 | 2.20 | 2.14 | 2.08 | 2.27 |
| >50% LPG + wood |  | 0.40 | 0.20 | 12.06 | 0.05 | 0.39 | 0.43 | 0.40 | 0.33 | 0.53 |
| 100% LPG |  | 0.00 | . | . |  | 0.00 | 0.00 | 0.00 | 0.00 | 0.00 |
| Kitchen type |  |  |  |  |  |  |  |  |  |  |
| Indoor |  | -0.17 | 0.43 | -0.45 | 0.69 | 0.12 | -0.01 | -0.21 | -0.20 | -0.34 |
| Temporary hut |  | 1.13 | 0.48 | 2.43 | 0.02 | 1.22 | 1.15 | 1.16 | 1.24 | 0.86 |
| Outdoor |  | 0.00 | . | . |  | 0.00 | 0.00 | 0.00 | 0.00 | 0.00 |
| Functional chimney |  |  |  |  |  |  |  |  |  |  |
| No |  | 0.06 | 0.17 | 0.35 | 0.72 |  |  |  |  |  |
| Yes |  | 0 |  |  |  |  |  |  |  |  |
| Adjusted R^2^ = 0.57; P-value for F-statistics <0.0001 | | | | | | | | | | |
